# A genome-wide CRISPR screen identifies ZCCHC14 as a host factor required for hepatitis B surface antigen production

**DOI:** 10.1101/718940

**Authors:** Anastasia Hyrina, Christopher Jones, Darlene Chen, Scott Clarkson, Nadire Cochran, Paul Feucht, Gregory Hoffman, Alicia Lindeman, Carsten Russ, Frederic Sigoillot, Kyoko Uehara, Lili Xie, Don Ganem, Meghan Holdorf

## Abstract

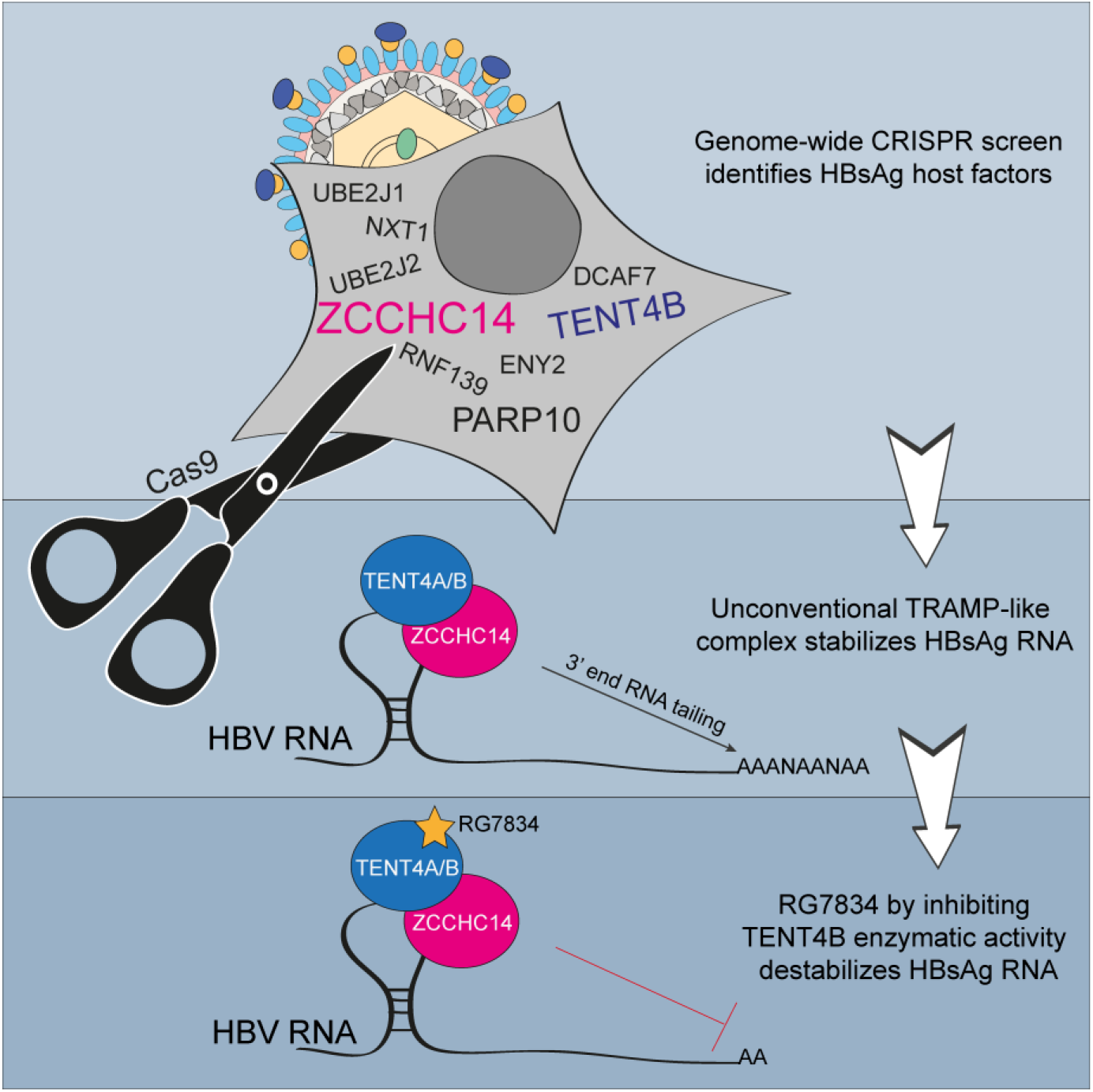

**SUMMARY:** A hallmark of chronic hepatitis B virus (CHB) infection is the presence of high circulating levels of non-infectious small lipid HBV surface antigen (HBsAg) vesicles. Although rare, sustained HBsAg loss is the idealized endpoint of any CHB therapy. A novel small molecule RG7834 has been previously reported to inhibit HBsAg expression by targeting terminal nucleotidyltransferase protein 4A and 4B (TENT4A and TENT4B). In this study, we describe a genome-wide CRISPR screen to identify other potential novel host factors required for HBsAg expression and to gain further insights into the mechanism of RG7834. We report more than 60 genes involved in regulating HBsAg and identified novel factors involved in RG7834 activity, including a zinc finger CCHC-type containing 14 (ZCCHC14) protein. We show that ZCCHC14, together with TENT4A/B, stabilizes HBsAg expression through HBV RNA tailing, providing a potential new therapeutic target to achieve functional cure in CHB patients.

## INTRODUCTION

Chronic hepatitis B virus (CHB) infection is a global public health concern affecting approximately 257 million people leading to risk of severe liver disease, including cirrhosis and hepatocellular carcinoma (Trepo et al., 2014). CHB is attributed to the maintenance of an intrahepatic pool of the viral covalently closed circular DNA (cccDNA), which serves as the transcriptional template for all viral gene products required for replication (Nassal, 2015). The most widely used antiviral therapies, nucleos(t)ide analogs, are effective at preventing infectious virus production and spread, but have no impact on cccDNA or the expression of HBsAg and other viral gene products. During CHB, HBsAg is expressed in large excess and circulates within small, non-infectious lipid vesicles which have been suggested to interfere with the host immune responses ((Lebosse et al., 2017) and reviewed in (Bertoletti and Ferrari, 2012; Faure-Dupuy et al., 2017)). Although rarely seen in response to existing drugs, loss of circulating HBsAg is clinically correlated with viral clearance. Along with seroconversion, sustained HBsAg reduction is considered an important clinical endpoint for HBV therapies and a key parameter associated with functional cure (Zoulim and Durantel, 2015).

To investigate mechanisms of reducing HBsAg, we performed a genome-wide CRISPR screen resulting in the identification of several novel host factors required for HBsAg expression. Using this approach, we also interrogated the mechanism of a small molecule inhibitor of HBsAg expression (RG7834), which was recently shown to destabilize HBV transcripts through interactions with the non-canonical poly(A) RNA polymerases TENT4B (PAPD5) and TENT4A (PAPD7) (Han et al., 2018; Mueller et al., 2019; Mueller et al., 2018). Here we provide strong evidence that a newly implicated host factor, ZCCHC14, together with TENT4B and TENT4A, stabilizes HBsAg expression through HBV RNA tailing (nontemplated addition to the 3’ terminus of the RNA). This mechanism is dependent on the HBV post-transcriptional regulatory element (PRE) and is disrupted by RG7834. These results not only further expand our understanding of how HBsAg expression is regulated, but also provide an avenue for potential new therapeutic targets for drug development.

## RESULTS

### A genome-wide CRISPR screen to identify host factors required for HBV transcription

To identify host factors required for HBsAg expression, we established a FACS-based genome-wide CRISPR screen in a HepG2 cell line containing an integrated 1.1-mer HBV genome (HepG2-HBV) (Figure 1A). Genome editing by CRISPR was enabled through the transduction of HepG2-HBV cells with lentivirus expressing Cas9 and a clonal cell line was selected that maintained stable high-level Cas9 expression over several passages (Cas9 HepG2-HBV, Figure S1A). The efficiency of genome editing was evaluated using a FLAER assay that measures disruption of the cellular enzyme phosphatidylinositol glycan complementation class A (PIG-A) in glycosylphosphatidylinositol (GPI)-anchor protein biosynthesis pathway (Brodsky, 2009; Estoppey et al., 2017). Using this method, Cas9 editing efficiency at 21 days was determined to be >90% (Figure S1B) and results were confirmed by next-generation sequencing (NGS) across the PIG-A gene locus (Figure S1C). Editing efficiency of the HBsAg gene in the integrated HBV genome was also evaluated by FACS analysis of intracellular HBsAg and an AlphaLISA assay of extracellular HBsAg at 14 and 21 days post-transduction. Intracellular HBsAg levels were reduced by 83-91% and extracellular HBsAg levels were reduced by 47-90%, depending on the sgRNA (Figures S1D and S1E). High efficiency editing of the integrated HBV genome was confirmed by NGS analysis (Figure S1F).

**Figure 1.**
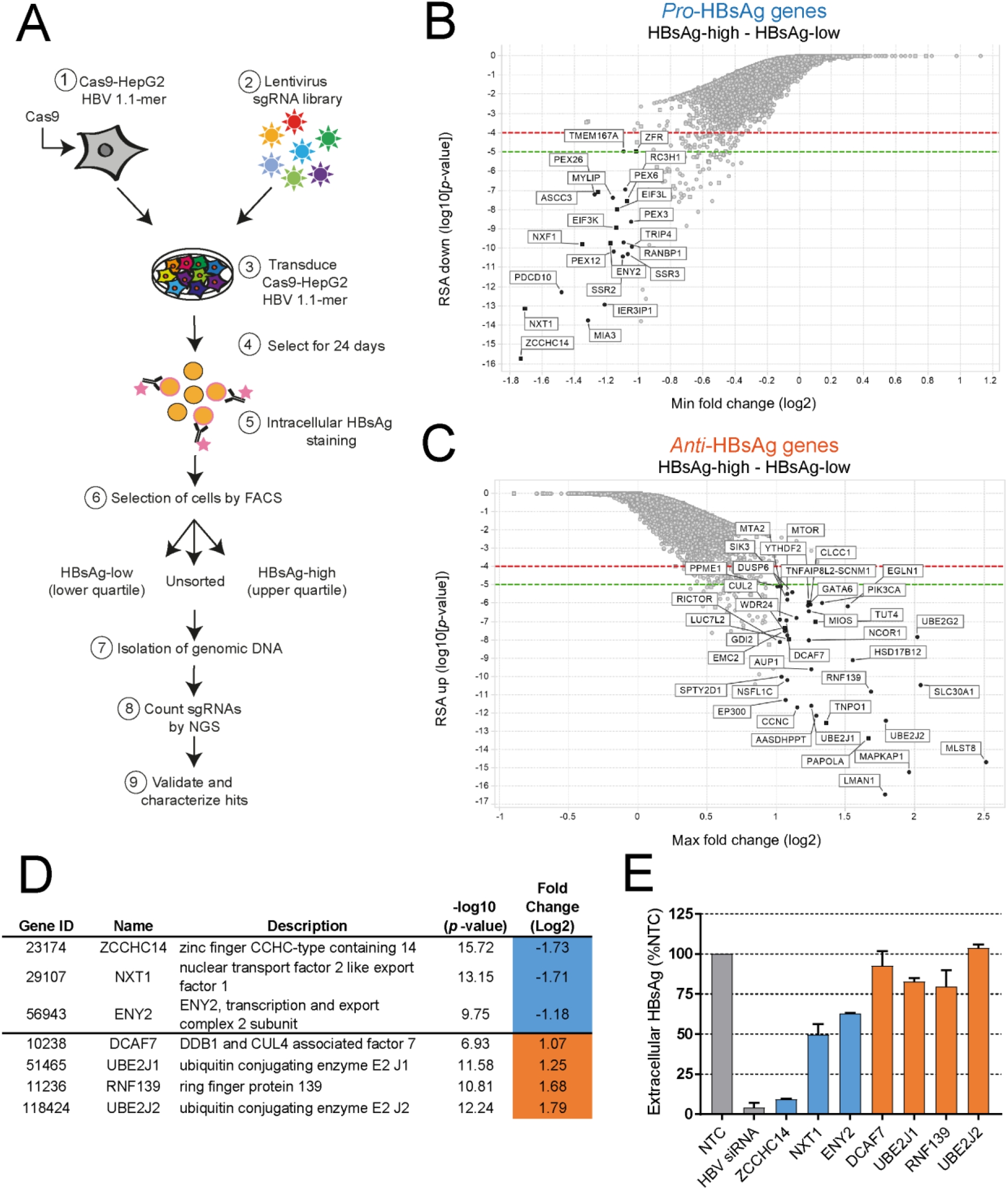
Novel host factors involved in HBsAg regulation identified by genome-wide CRISPR screen. **(A)** Workflow of the CRISPR screening process. **(B**-**C)** Waterfall plots showing potential genes involved in HBsAg expression (“pro-HBsAg”) **(B)**, and genes that act as HBsAg restriction factors (“anti-HBsAg”) **(C)** using a cut-off of at least 2-fold change and RSA < −5. Statistical significance was quantified using RSA analysis of two biological replicates. **(D)** A summary of pro-HBsAg (blue) and anti-HBsAg (orange) genes validated by siRNA-mediated knockdown of individual genes. RSA and fold change values are from the CRISPR screen. **(E)** Extracellular HBsAg levels analyzed by AlphaLisa following siRNA-mediated knockdown of the putative pro-HBsAg and anti-HBsAg genes in HepG2-HBV for 5 days. The graph shows mean values ± standard deviation (SD) of two biological replicates.

To perform the genome-wide CRISPR screen, Cas9 HepG2-HBV cells were transduced with a pooled puromycin-selectable lentiviral sgRNA library targeting 19,050 human protein-coding genes with five sgRNAs per target gene (DeJesus et al., 2016). The general workflow is shown in Figure 1A. Briefly, Cas9 HepG2-HBV cells were transduced with lentivirus at low multiplicity of infection (MOI = 0.5) in two independent replicates, followed by puromycin selection for 24 days to allow efficient CRISPR gene editing. Cells were then stained using an HBsAg antibody and sorted by FACS into HBsAg-high and HBsAg-low populations (Figure S2A). Genomic DNA was then isolated and analyzed by NGS to determine the abundance of each sgRNA-encoding sequence within the sorted populations.

With the exception of one sample (RE-04-UV84, unsorted replicate 2), which was removed from downstream analysis, all samples had a minimum of 1×10^7^ NGS reads (Figure S2B); approximately 90% of which aligned to expected barcodes from the pooled libraries (Figure S2C). Genes were then assigned a score based on the log_2_-transformed fold-change of sgRNA counts in unsorted samples compared to the sgRNA counts in the input CRISPR library. sgRNAs targeting essential cellular factors, including the ribosomal subunits (RPS8, RPS3A, and RPS13), were found in lower abundance than in the input CRISPR library (Figure S2D), while sgRNAs against the tumor suppressor gene TP53 were highly enriched (Figure S2E). Observation of the expected behavior of these sgRNAs validated the pooled CRISPR genome-wide screening approach.

### Novel host factors involved in HBsAg production identified in genome-wide CRISPR screen

To identify host factors involved in HBsAg expression, enrichment of sgRNA-targeted genes was compared between populations expressing high and low levels of HBsAg. Using a 2-fold cutoff, this analysis identified 22 genes that appeared to promote HBsAg expression (“pro-HBsAg”) and 38 genes that reduced HBsAg levels (“anti-HBsAg”) (Figure 1B and 1C). Gene ontology analysis revealed significant enrichment of peroxisomal pathway(s) factors among genes that promoted HBsAg expression (redundant siRNA activity (RSA) = 17.8, with enrichment in 14 genes), while anti-HBsAg genes included those from the ER-associated protein degradation (ERAD) pathway (RSA = 4.72, with enrichment in 5 genes) (Figures S3A and S3B, Table S1). The most enriched pro-HBsAg gene was zinc finger CCHC-type containing 14 (ZCCHC14). ZCCHC14 is a 949 amino acid protein containing one putative zinc-binding motif of the form Cys-X2-Cys-X4-His-X4-Cys, referred to as a ‘Zn-knuckle’ (Fields et al., 2008).

To validate the impact of selected host factors on HBV replication, siRNA mediated knockdown of individual genes was performed (Figure 1D). HepG2-HBV cells were transfected with siRNA SMARTpools targeting each individual gene and HBsAg levels were determined by FACS (intracellular) and HBsAg AlphaLISA (extracellular) assays at 5 days post-transfection. Gene knockdown was confirmed by qRT-PCR (Figure S4A). Consistent with the screening results, knockdown of pro-HBsAg genes ZCCHC14, NXT1, or ENY2 reduced both intracellular and extracellular HBsAg levels, with knockdown of ZCCHC14 leading to the most pronounced decrease in HBsAg production (Figure 1E and Figure S4B). Knockdown of the anti-HBsAg genes DCAF7, UBE2J1, RNF139, or UBE2J2 boosted intracellular HBsAg levels relative to NTC siRNA (Figure S4B). Extracellular HBsAg levels, however, were not changed (Figure 1E) suggesting that the cellular machinery has reached maximum secretion of HBsAg (Huovila et al., 1992). Together, these results validate selected genes as host factors that either promote or restrict HBsAg expression.

### ZCCHC14, TENT4B, and PARP10 are involved in RG7834 activity

To further explore the factors that regulate HBsAg expression, we investigated a small molecule inhibitor of HBV replication, RG7834 (EC_50_ = 0.4nM, Figure S4C) and expanded the genome-wide CRISPR screen to include compound-treated cells. Cas9 HepG2-HBV cells were transduced with the pooled puromycin-selectable lentiviral sgRNA library, followed by puromycin selection for 14 days. Transduced cells were then divided into three groups and treated with RG7834 (30pM or 30nM) or DMSO for 10 days. A low dose (30pM; ~IC_20_) of RG7834 was used to identify factors (when knocked out) could potentiate the activity of the compound; a higher dose of RG7834 (30nM; ~IC_80_) was used to identify factors required for RG7834 activity.

Knockout of poly(ADP)-ribose polymerase family member 10 (PARP10), a protein known to promote tumorigenesis by alleviating replication stress (Schleicher et al., 2018) possibly potentiates RG7834 activity (Figure 2A). This finding was validated by treating HepG2-HBV cells with OUL35, a selective small-molecule inhibitor of PARP10 (Venkannagari et al., 2016). When cells were treated with OUL35 (1μM) alone for 5 days, no effect was detected in extracellular HBsAg levels. However, when cells were treated with both OUL35 (1μM) and RG7834 (1nM), extracellular HBsAg levels were significantly lower compared to RG7834 treatment alone (Figure 2B) without causing cellular toxicity (data not shown).

**Figure 2.**
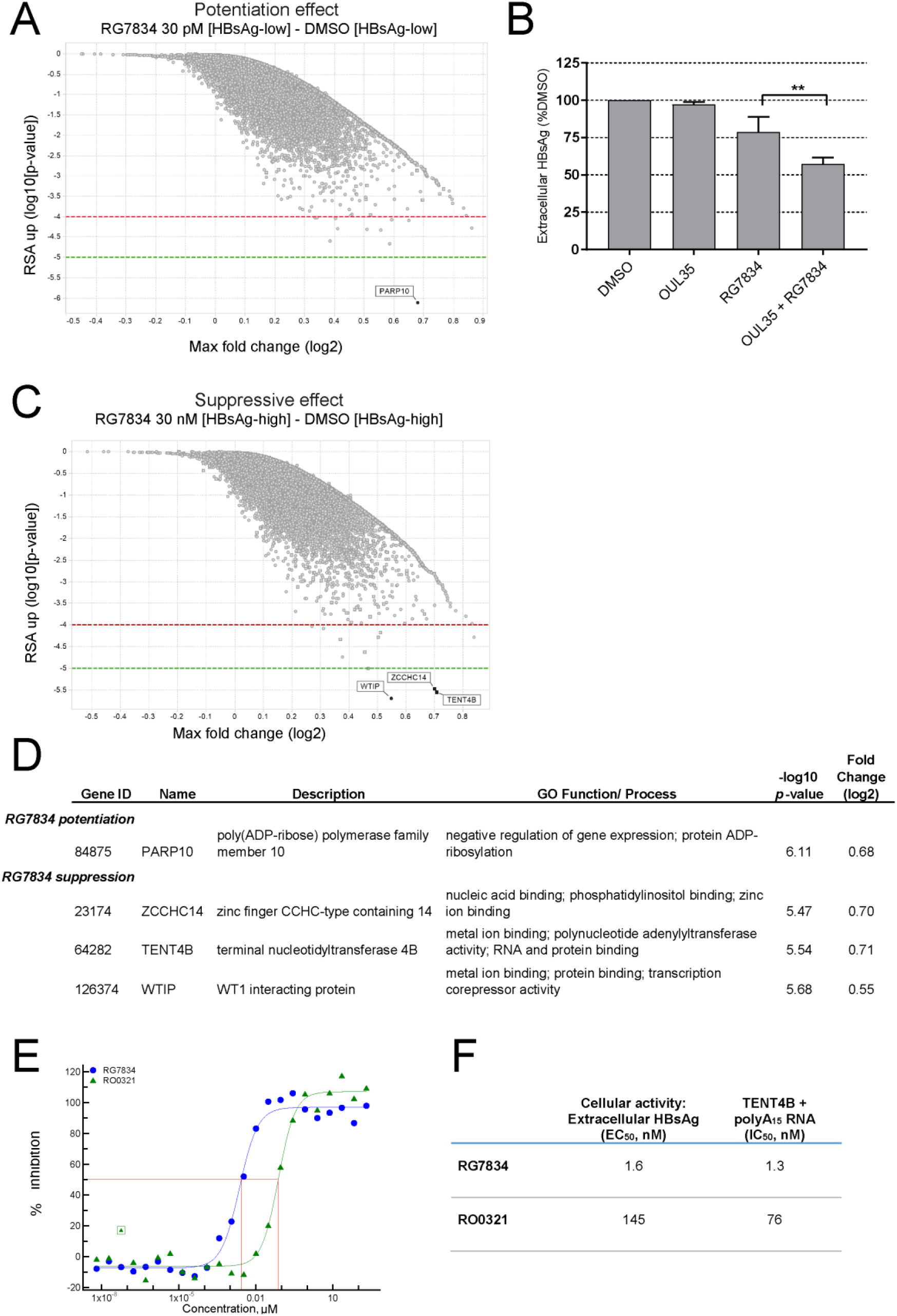
Genome-wide CRISPR screen identifies efficacy targets of RG7834. **(A)** Waterfall plot of genes (when knocked out) have RG7834 potentiation. **(B)** AlphaLISA analysis of extracellular HBsAg in HepG2-HBV cells treated with OUL35 (1μM) ± RG7834 (1nM) or DMSO control for 5 days. The graph shows mean values ± SD of two biological replicates. Statistical analysis was performed using ordinary one-way ANOVA in GraphPad Prism 7.04. **, represents p-value ≤ 0.01. **(C)** Waterfall plot of genes essential for activity of RG7834. **(D)** A summary of top gene hits from RG7834 potentiation and suppression analyses including individual gene ontology (GO) function and process analysis. Candidate gene hits were selected with the stringent threshold of RSA < −5.0. **(E)** Dose response curve of polyadenylation activity of purified TENT4B on a template RNA (polyA_15_) in the presence of RG7834 or RO0321. **(F)** Summary of RG7834 and RO0321 IC50 and EC50 values based on at least two independent experiments.

Analysis of host genes essential for activity of RG7834 identified TENT4B, WT1 interacting protein (WTIP), and ZCCHC14 (Figure 2C and 2D). WTIP showed the lowest fold-change among these genes and was not examined further. TENT4B has been previously identified as a target of RG7834 from a yeast-three-hybrid approach, but had not been validated biochemically (Mueller et al., 2019). We therefore established an assay to measure the polymerase/polyadenylation activity of purified TENT4B. In this assay, polyadenylation of a template RNA (polyA15) was monitored indirectly by measuring ATP depletion over time using Kinase-glo (Promega). To rule out non-specific ATP hydrolysis, we also measured AMP/ADP production by ADP-glo (Promega). Purified TENT4B demonstrated robust RNA-dependent ATP depletion, without detectable levels of free AMP/ADP (Figure S4D). Addition of RG7834 was found to inhibit TENT4B polymerase activity with an IC_50_ = 1.3nM, whereas an RG7834 enantiomer (RO0321) was ~60-fold less potent (Figure 2E and 2F). These findings suggest that RG7834 is an inhibitor of TENT4B polymerase/polyadenylation enzymatic activity.

### ZCCHC14 works in conjunction with the TENT4A/B to tail HBV RNAs

In vivo, TENT4B functions as part of the larger TRAMP-like complex (Hamill et al., 2010; Houseley and Tollervey, 2009). The complex consists of three subunits highly conserved across eukaryotic organisms: a non-canonical poly(A) polymerase (TENT4B or TENT4A in mammals; Trf4/5 in yeast), a Zn-knuckle protein (ZCCHC7 in mammals; Air1/2 in yeast), and an RNA helicase (MTR4) (Fasken et al., 2011; Hamill et al., 2010). Very little has been reported regarding ZCCHC14, except that it is also a Zn-knuckle protein. To evaluate the roles of ZCCHC14 and multiple members of the TRAMP-like complex in HBV replication and gene expression, HepG2-HBV cells were transfected with siRNAs for 5 days and gene knockdown confirmed by qRT-PCR (Figure S4A). Knockdown of TENT4B, TENT4A, or ZCCHC7 alone had minimal effect on intracellular or extracellular HBsAg protein levels. Simultaneous knockdown of TENT4A/B was required for a moderate reduction in intracellular HBsAg and a pronounced reduction in extracellular HBsAg (Figures 3A and 3B). In contrast, knockdown of ZCCHC14 alone strongly suppressed HBsAg to levels comparable to RG7834 treated cells with minimal impact on cell viability (Figures 3A-D). Knockdown of both ZCCHC14 and TENT4B had the same effect on intracellular and extracellular HBsAg as knockdown of ZCCHC14 alone (data not shown). ZCCHC14 does not appear to control TENT4B expression, since depletion of ZCCHC14 did not alter TENT4B protein levels (Figure S4E). We reconfirmed these findings and the role of ZCCHC14 and the TRAMP-like members in HBsAg expression using HBV de novo infected HepG2-NTCP1 cells (Figures S4F and S4G). We note that siRNA-mediated knockdown of ZCCHC14 (as well as TENT4A/B) also reduced intracellular levels of HBV DNA (Figure 3C) – an effect that would not be expected from modulation of HBsAg mRNA levels alone. This suggests that the effect on RNA accumulation may extend to pre-genomic RNA as well.

**Figure 3.**
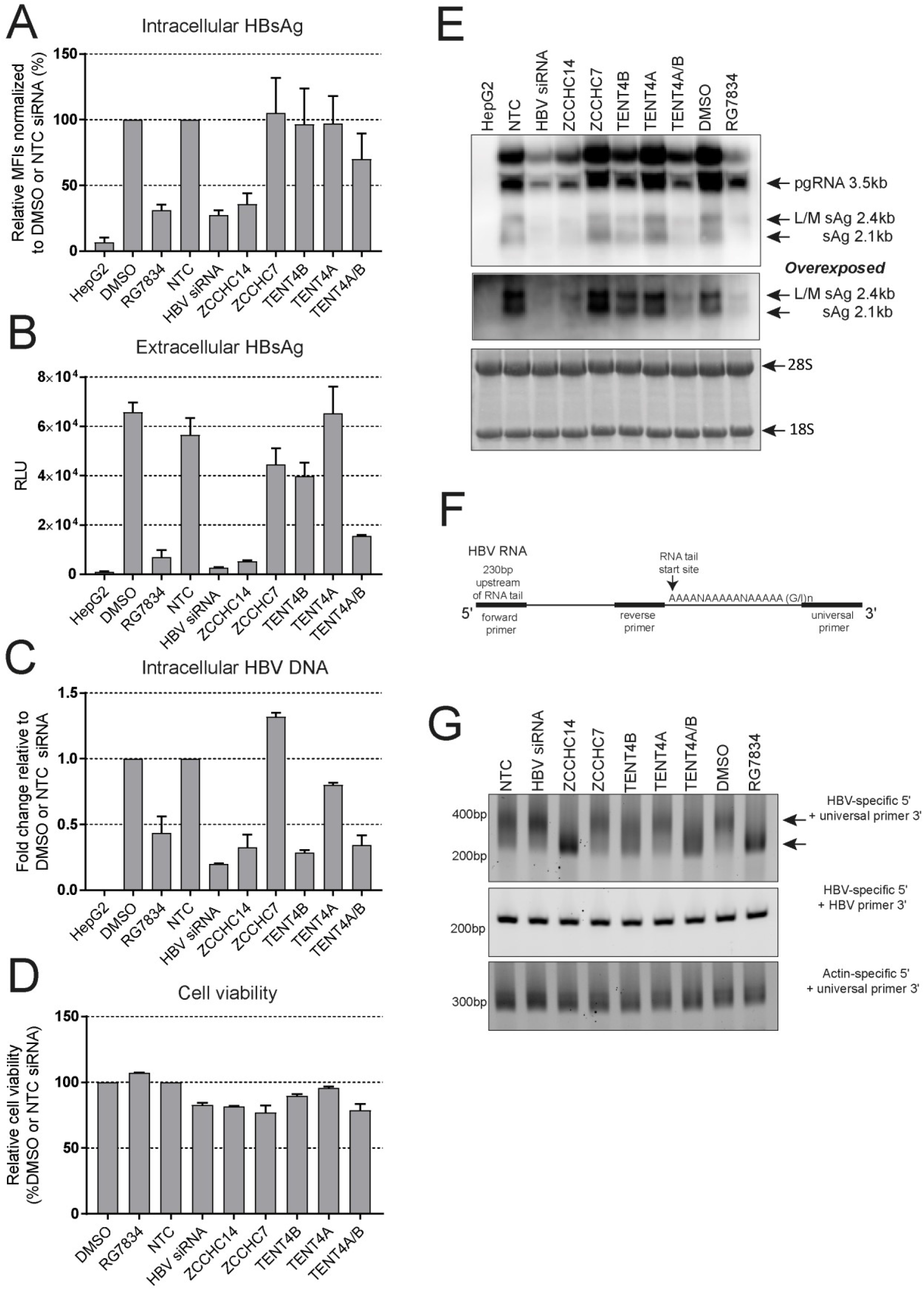
ZCCHC14 works in conjunction with the TENT4A/B to affect HBV RNA tailing. HepG2-HBV cells were treated with RG7834 (1μM) or transfected with ON-TARGET *plus* siRNA SMARTpools and controls. (A) FACS analysis of intracellular HBsAg levels; (B) AlphaLISA analysis of extracellular HBsAg levels, (C) qPCR analysis of total intracellular HBV DNA; and (D) Cell Titer Glo analysis of cell viability 5 days post-treatment. The graphs show mean values ± SD of at least two biological replicates. **(E)** Northern blotting analysis of HBV RNA transcripts. **(F)** Schematic demonstration of primers design for RNA tail assay. **(G)** PCR amplification results following the RNA tail assay. The top arrow corresponds to the average RNA tail length observed in HBV transcripts. The bottom arrow corresponds to the shortened length observed upon certain treatments.

One of the key factors involved in mRNA stability regulation by the TRAMP complex is the length of the polyA tail (Vanacova et al., 2005). Consistent with previous reports (Mueller et al., 2019), we saw a decrease in RNA abundance and moderate increases in electrophoretic mobility of all HBV transcripts (including pre-genomic RNA) isolated from cells treated with RG7834, TENT4B siRNA, or TENT4A/B siRNAs; knockdown of either TENT4A or ZCCHC7 had no observable impact on HBV RNA mobility (Figure 3E). To evaluate the tail lengths of HBV transcripts, we performed an RNA tail length assay (Figure 3F) and using HBV-specific forward and reverse primers positioned upstream of the RNA tailing site verified target-specific amplification. In this assay, RG7834 treatment and ZCCHC14 knockdown had the most profound effect on HBV RNA tails length, with ZCCHC14 knockdown closely phenocopying RG7834 inhibitor treatment. TENT4B and TENT4A/B knockdown also reduced HBV RNA tails length, but to a lesser extent (Figure 3G). Actin RNA tails were unchanged, indicating that the effect was not due to a global reduction in RNA tailing.

### Intact stem loop alpha (SLα) of HBV PRE is critical for ZCCHC14 activity

The activity of RG7834 has previously been shown to depend on a stem loop structure within the 5’ end of the PRE, termed SLα. Deletions and mutations of SLα abrogated the inhibitory activity of RG7834, although the mechanism of this effect is still unclear (Zhou et al., 2018). To determine whether this RNA element works in concert with the putative unconventional TRAMP-like complex, we generated a non-infectious HBV reporter cell line (HBV Rluc) in which the entire HBsAg coding region was replaced with a Renilla luciferase (Rluc) reporter downstream of the preS1/S2 promoters, while retaining the entire PRE and authentic HBV 3’UTR (Figure 4A). To evaluate the critical residues of the PRE SLα, mutations within the SLα region (ntd 1292-1321) were generated (HBV SLα 4x mut). A HepG2 cell line expressing Rluc from a stably integrated CMV plasmid served as a control (CMV Rluc).

**Figure 4.**
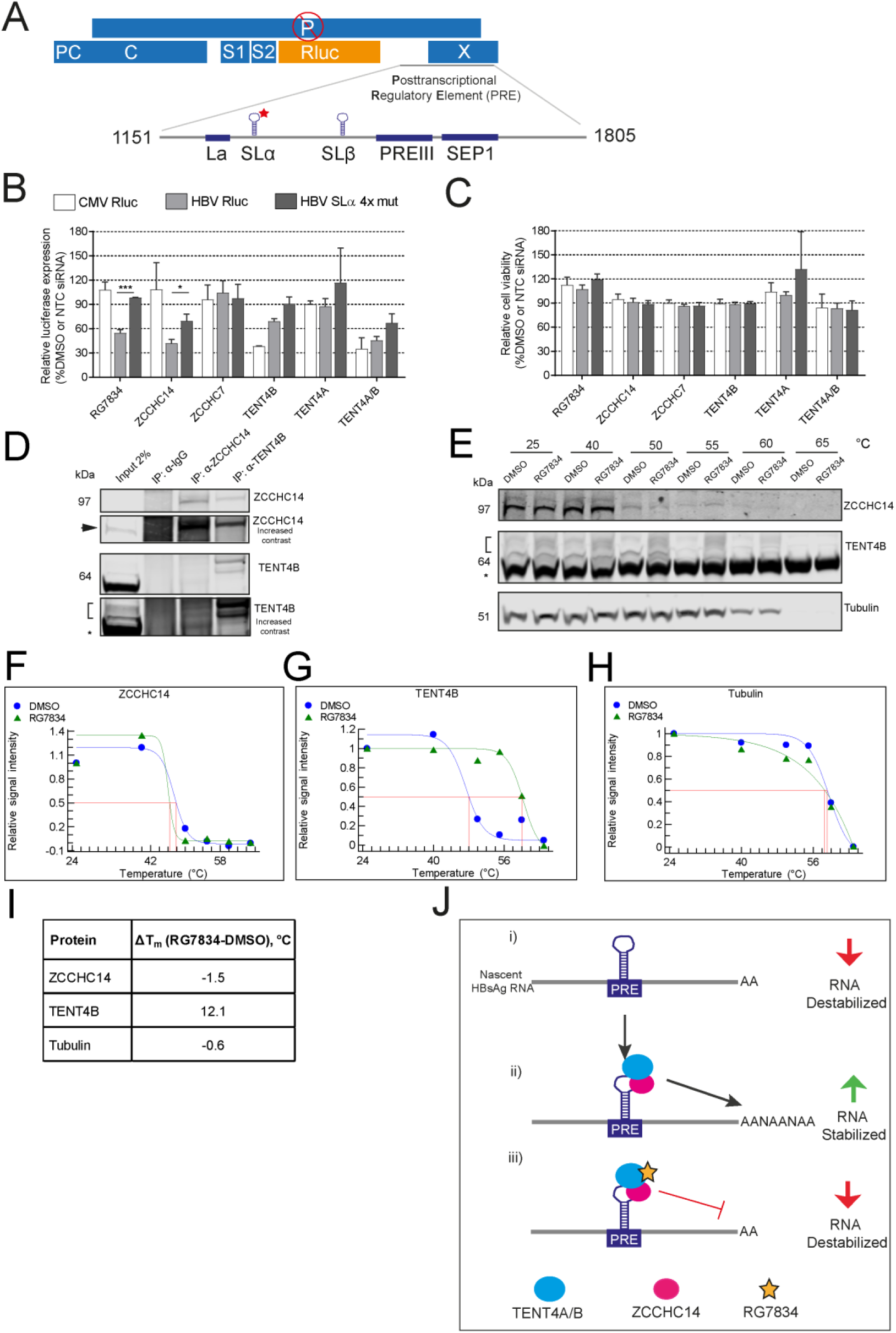
Intact SLα of HBV PRE is critical for ZCCHC14 activity in complex with TENT4B. **(A)** HBV Rluc and the HBV SLα 4x mutant construction. **(B-C)** Reporter cell lines were treated with RG7834 (1μM) or transfected with ON-TARGET *plus* siRNA SMARTpools monitoring luciferase activity as a proxy for HBsAg expression **(B)** and cell viability by Cell Titer Glo (Promega) **(C)** 3 days post-treatment. The graphs show mean values ± SD of three biological replicates. Statistical analysis was performed using ordinary one-way ANOVA. *, p-value ≤ 0.05; ***, ≤ 0.001. **(D)** Analysis of ZCCHC14 and TENT4B interaction by co-IP. Proteins from the input (before IP) and the pulldowns were analyzed by western blotting. A bracket marks several TENT4B isoforms. *, in TENT4B blots indicates a non-specific band. **(E)** Cellular thermal shift assay to investigate RG7834 binding to cellular proteins. Protein expression levels were determined by western blotting and quantified by densitometry. **(F-H)** Relative signal intensities were fit using Boltzman model in XLfit. **(I)** ΔT_m_ was quantified based on T_m_ of the proteins in the presence of RG7834 or DMSO. **(J)** Proposed model of a novel complex consisting of ZCCHC14 and TENT4A/B in stabilization of HBV RNAs through RNA tailing.

The luciferase signal and cell viability of the reporter HepG2 cell lines were evaluated three days post treatment with either RG7834 or siRNA treatment (Figures 4B and 4C). As expected, RG7834 treatment decreased luciferase expression by the HBV Rluc reporter cell line, with inhibitory activity lost in the presence of HBV SLα mutations. Similarly, knockdown of ZCCHC14 specifically reduced luciferase activity for HBV Rluc, and this inhibition was less pronounced for the SLα mutant. In addition, knockdown of TENT4B and the TENT4A/B reduced luciferase activity in both HBV Rluc and in CMV Rluc, indicating that this complex can regulate expression of diverse genes. The HBV SLα mutation slightly diminished the effects of both TENT4B and TENT4A/B knockdown. Neither knockdown of ZCCHC7 nor TENT4A had an effect on luciferase activity in any of the cell lines. Our results indicate that ZCCHC14 plays a role specific to HBV, whereas TENT4A/B may be regulating a broader range of RNAs. These findings suggest that ZCCHC14, possibly in complex with TENT4A/B, functions to promote HBsAg expression through the SLα within the PRE region of HBV.

Given these findings, we next sought to further characterize this potential complex between ZCCHC14 and TENT4B. Co-immunoprecipitation using a TENT4B antibody in HepG2-HBV cells was able to pull-down ZCCHC14 (Figure 4D). We were unable however, to confirm this interaction using a ZCCHC14 antibody to pulldown TENT4B. This could be due to low levels of endogenous TENT4B isoforms. To further investigate the binding of RG7834, we used a cellular protein thermal shift assay to determine the effect of RG7834 binding on the thermal stability of TENT4B and ZCCHC14 in cellular lysates. In this assay, compound binding is predicted to increase the thermal stability of its target protein. The results demonstrated that RG7834 increased the thermal stability of TENT4B (∆Tm = 12°C), but not ZCCHC14 (Figure 4E-I).

## DISCUSSION

One of the strongest confirmed pro-HBsAg factors identified from the genome-wide CRISPR screen was ZCCHC14. Interestingly, ZCCHC14 also came up as an efficacy target of RG7834, along with a previously identified target of the compound, TENT4B (Mueller et al., 2019). Our results suggest a novel complex consisting of ZCCHC14 and TENT4A/B, which functions to stabilize HBV mRNA by promoting 3’-end tailing and thereby likely preventing degradation of the tail by exonucleases (Figure 4J). This complex resembles the TRAMP-like machinery, but with ZCCHC14 replacing ZCCHC7. For cellular mRNAs, polyadenylation by the TRAMP-like complex was originally shown to target RNAs for degradation by the nuclear exosome (Boele et al., 2014; LaCava et al., 2005). Recently, however, TENT4A/B were shown to protect mRNAs from degradation by adding mixed poly(A) tails with intermittent non-adenosine residues (Lim et al., 2018). It was also noted that transcripts encoding secreted proteins are frequently guanylated and are affected strongly by TENT4A/B depletion. Our results suggest a protective activity for the modified TRAMP-like complex on HBsAg mRNA, although it remains to be shown if this involves non-adenosine addition.

HBV RNA tailing is unique due to the circular, overlapping nature of the cccDNA genome. The polyadenylation signal (UAUAAA) on the terminally redundant genomic transcript (3.5-kb) is known to function poorly, in fact it is recognized only upon the second encounter with the polyadenylation machinery (Simonsen and Levinson, 1983). HBV employs a number of RNA elements to boost polyadenylation and translation (Russnak and Ganem, 1990; Simonsen and Levinson, 1983), including a 3’-PRE that enhances viral gene expression and mRNA nuclear export (Huang and Liang, 1993; Huang and Yen, 1994). Zhou et al. recently reported that RG7834-mediated reduction of HBV transcripts requires the PRE, with SLα important for activity (Zhou et al., 2018). Here we show that the PRE is also critical for the functional activity of ZCCHC14 and that ZCCHC14 physically interacts with TENT4B. Based on these data, we propose a model in which ZCCHC14 recruits an unconventional TRAMP-like complex to HBV mRNA through interaction with the PRE sequence. Moreover, RG7834 by binding TENT4B and inhibiting its enzymatic activity within the TENT4A/B-ZCCHC14-PRE complex inhibits RNA tailing activity and results in HBV RNA destabilization (Figure 4J).

In addition to identifying genes essential for RG7834 activity, we identified PARP10 as a host factor that negatively impacts RG7834 potency. Indeed, a selective small molecule inhibitor of PARP10 enzymatic activity (Venkannagari et al., 2016) appears to potentiate the activity of RG7834. PARP10 is one of 17 diphteria toxin-like ADP-ribosyltransferases (ARTD)/PARP family enzymes that transfer single or poly-ADP-ribose groups to a target protein. This post-translational modification can have a wide range of effects, from modulating enzymatic activity to facilitating ubiquitination and subsequent degradation of targeted proteins (Kuny and Sullivan, 2016). ARTD family members have strong connections to host–pathogen interactions, showing both pro- and anti-viral activities, including HBV integration into host DNA (Dandri et al., 2002; Gao et al., 2002; Kuny and Sullivan, 2016).

Using a genome-wide CRISPR screen, we identified 60 genes that appeared to promote or reduce HBsAg levels. Unexpectedly, we identified several ubiquitination proteins with anti-HBsAg activity. Disruption of UBE2J1, RNF139, UBE2J2 and DCAF7 resulted in increased HBsAg levels, suggesting HBsAg could be a natural substrate for therapeutic modulation by targeted protein degradation approach (Lai and Crews, 2017). Also of interest was the peroxisomal pathway. It was striking that 14 peroxisome-associated genes were enriched in the pro-HBsAg set. Some of these encode structural components of the peroxisome, while others encode genes that affect targeting of proteins to this organelle. We do not know the basis for the strong association of peroxisome-associated genes with HBsAg biogenesis but it has previously been shown that HBsAg biosynthesis involves budding of nascent chains into a pre-Golgi compartment (Huovila et al., 1992; Simon et al., 1988). Peroxisomes are also known to derive from the ER; our results could indicate that the compartment in which HBsAg forms may share components with the peroxisome or a precursor thereof. Also, the role of peroxisomes in lipid metabolism, including the beta-oxidation of fatty acids and the biosynthesis of phospholipids maybe important for HBsAg assembly into lipoprotein particles (Lodhi and Semenkovich, 2014; Satoh et al., 1990). Several studies have shown that peroxisome proliferator-activated receptor alpha agonists increase transcription and replication of HBV (Guidotti et al., 1999; Tang and McLachlan, 2001).

In conclusion, a genome-wide CRISPR screen provided a powerful strategy for identifying new genes and pathways that control HBsAg expression. In addition, we demonstrate this unbiased approach as a tool for dissecting the potential mechanism of inhibitors of HBsAg expression. In particular, we provide the first evidence for a role of ZCCHC14 in augmenting the stability of HBsAg mRNA through HBV RNA tailing by TENT4A/B as part of an unconventional TRAMP-like complex. These results suggest a potential new therapeutic target for interrupting HBV replication and gene expression in CHB.

## ACKNOWLEDGEMENTS

We thank Tiffany Tsang for her help with the protein reagents and Catherine Jones for useful suggestions and editing of the manuscript.

## AUTHOR CONTRIBUTIONS

A.H., C.J., D.G., G.H. and M.H. conceived and designed the study. A.H. performed the experiments. D.C. generated the HBV reporter cell lines. N.C. performed FACS-sorting and genomic DNA isolation of samples during the genome-wide CRISPR screen. S.C., A.L. and C.R. performed the NGS. F.S. analyzed the sgRNA distribution. P.F. performed TENT4B ATP depletion assay. K.U. purified TENT4B. L.X. provided protein sciences support. A.H., C.J., and M.H. wrote the manuscript. All authors approved the manuscript.

## DECLARATION OF INTERESTS

All the authors are employees of Novartis Institutes for BioMedical Research.

## STAR METHODS

### Mammalian cell culture, cell line generation and small molecule

HepG2 cells (obtained from ATCC) were cultured in DMEM/F12, GlutaMAX (Life Technologies), 10% fetal bovine serum (FBS, Gibco), and 100 U/mL penicillin/streptomycin (Gibco) at 37°C and 5% CO_2_. HepG2-hNTCP and HepG2-HBV were generated and cultured as previously described (Tropberger et al., 2015). The Cas9 HepG2-HBV cell line was generated through lentiviral delivery of Cas9 followed by selection with blasticidin (Life Technologies) and isolation of single cell clones (DeJesus et al., 2016). Individual clones were tested for Cas9 expression by flow-cytometry on mouse Cas9 PE conjugated (7A9-3A3, Cell Signaling #35193) stained cells at a 1:200 dilution. The Cas9+ clone 16 was validated by on-target NGS using the VarDict variant calling algorithm (Lai et al., 2016), FLAER described in (Estoppey et al., 2017) and gRNAs targeting HBV genome (gRNA1: TAACTGAAAGCCAAACAGTG; gRNA2: CAAAAGAAAATTGGTAACAG; gRNA3: CCTGCTGCGAGCAAAACAAG).

The non-infectious reporter cell lines (HBV Rluc, HBV SLα 4x mut, and CMV Rluc) were generated in HepG2 cells by transfection and hygromycin (Corning) selection of the corresponding plasmids. For the HBV Rluc cell line, the Rluc reporter was inserted in HBV genotype B genome under internal S1/S2 promoters disrupting P polymerase. For the 4x SLα mutant, four mutations were introduced into the SLα region (ntd 1292-1321) of the HBV Rluc plasmid as previously described in (Zhou et al., 2018).

RG7834 and RO0321 were synthesized at Novartis based on the reported structures (Han et al., 2018; Mueller et al., 2019; Mueller et al., 2018). The EC50 value for RG7834 on intracellular HBsAg in Cas9 HepG2-HBV cell line was determined by a 16-point dose-response ranging from 0-10 μM. The cells were treated for 10 days, replenishing media and compound every 3-4 days. Intracellular HBsAg levels were measured by FACS and a dose-response curve was plotted and analyzed with XLFit using dose response one site four parameter logistic model [204: fit = (A+((B-A)/(1+(10^((C-x)*D)))))] based on the samples’ median fluorescent intensities.

### Pooled genome-wide CRISPR screening

The human genome-wide sgRNA library targeting 19,050 protein-coding genes was designed, constructed, and packaged as described in detail (Potting et al., 2018). The screen was performed in duplicate by transduction of the sgRNA libraries split between two pools (Cpool1 and Cpool3) at an MOI determined by a 12-point dose-response ranging from 0 to 400 μL of viral supernatants in the presence of 5 μg/mL polybrene. Infection rate was determined by FACS as a percentage of red fluorescent protein (RFP)-positive cells and selection was optimized by determining the puromycin dose required to achieve >95% cell killing in 5 days.

For the genome-wide screen, cells were seeded at 1×10^8^ cells per 5-layered cell stack (Corning) in media containing 5 μg/mL polybrene and lentivirus at an MOI of 0.5. Twenty-four hours after infection, virus-containing media was replaced with fresh media containing puromycin for selection. Seven days after infection, cells were trypsinized and plated into new cell stacks at 1×10^8^ cells per cell stack; an aliquot of cells was analyzed by FACS to confirm infection and selection efficiency. Cells were maintained in culture and split as needed to ensure confluence did not exceed 90%. Starting at day 15 after infection, cells were cultured in media containing either DMSO, RG7834 at low (30pM; ~IC20), or RG7834 at high (30nM; ~IC80) concentrations (Schirle and Jenkins, 2016) for an additional 10 days. For FACS analysis of the sgRNA libraries, 1×10^9^ cells per each condition were fixed with BD Cytofix/Cytoperm Fixation/Permeabilization Solution (BD Biosciences) for 20 min at 4°C, followed by staining with primary human HBsAg antibodies (CR8097; US9512201 at 1.5 μg/mL) for 1h at 4°C. Finally, cells were stained with APC-conjugated secondary antibody (Southern Biotech; 1:500) for 1h at 4°C. Single, RFP positive cells were sorted (BD ARIA Fusion) from the lower HBsAg quartile (HBsAg-low) or from the upper HBsAg quartile (HBsAg-high) with each replicate containing a minimum of 1×10^8^ cells per population. One hundred million unsorted cells were also collected as an input sample.

Illumina library construction, sequencing, and genome-wide CRISPR screen data analysis were performed as previously described (Potting et al., 2018). In brief, fixed cells were treated in de-crosslinking buffer overnight at 65°C (TNES: 10mM Tris-Cl pH 8.0, 100mM NaCl, 1mM EDTA, 1%SDS), and genomic DNA extracted using phenol:chloroform:isoamyl alcohol (Sigma). Illumina sequencing libraries were generated using 24 × 4μg PCR reactions per transduced sample with primers specific to the genome integrated lentiviral vector backbone sequence. PCR primer sequences have been previously reported (DeJesus et al., 2016). The resulting Illumina libraries were purified using 1.8×SPRI AMPure XL beads (Beckman Coulter) following the manufacturer’s recommendations and qPCR quantified using primers specific to the Illumina sequences following standard methods. Illumina sequencing libraries were pooled and sequenced on the HiSeq 2500 instrument (Illumina) with custom sequencing primers (DeJesus et al., 2016).

For genome-wide CRISPR screen data analysis, raw sequencing reads were aligned to the appropriate library through Bowtie analysis with no mismatches (Langmead et al., 2009). Differential fold-change estimates between samples were generated for each sgRNA using DESeq2 (Love et al., 2014) and effects on proliferation were assessed by comparison of the unsorted cell population to the input library. For gene-based hit calling, consistency of all sgRNAs per gene were considered (Konig et al., 2007) and plotted against the second strongest sgRNA as a representative. For gene-set enrichment analyses, the gene-set database was compiled from multiple sources including Reactome, NCBI Biosystems, and Gene Ontology, and the hits were calculated using a hypergeometric overrepresentation test. Benjamini-Hochberg–corrected p-values were calculated for each gene set and combination of input gene list size.

### Validation of genes by siRNA knockdown

To evaluate activity of individual genes against HBV in HepG2-HBV, HepG2-hNTCP, and HBV reporter cell lines, cells were plated either in a 6-well or 96-well format in antibiotic-free medium. The following day transfection mixtures were prepared by diluting siRNAs (Dharmacon) and DharmaFECT1 transfection reagent (Dharmacon) in Opti-MEM (Gibco) following manufacturer’s instructions. For the HBV siRNA control, we used the siRNA sequence previously described and validated (Morrissey et al., 2005). The ON-TARGET plus human siRNA SMARTpools were purchased from Dharmacon: ZCCHC14 (23174; L-014086-01), NXT1 (29107; L-017194-00), ENY2 (56943; L-018808-01), DCAF7 (10238; L-019999-00), UBE2J2 (118424; L-008614-00), RNF139 (11236; L-006942-00), UBE2J1 (51465; L-007266-00), ZCCHC7 (84186; L-014804-01), TENT4B (64282; L-010011-00), TENT4A (11044; L-009807-00). For the non-target control, the ON-TARGET plus non-targeting pool (D-001810-10) was used. Transfection complexes were formed at room temperature and added to the plated cells for 24 hours. RG7834 or DMSO was also added to the cells at day 1 post-plating. For de novo infection, HepG2-hNTCP cells were infected with cell culture-derived HBV two days post plating as previously described (Tropberger et al., 2015). Infected cells were incubated for an additional 7 days and HBsAg levels assessed by immunofluorescence. In HepG2-HBV cells, supernatants and cells were harvested and used for HBV marker analysis at 5-6 days post-transfection. In HBV reporter cells lines, luminescence was measured with Renilla-Glo Luciferase assay following manufacturer’s instructions at 3 days post-transfection. Cell viability was determined by Cell Titer Glo (Promega). All siRNA knockdowns were validated by qRT-PCR analysis of the targeted mRNAs. All experiments were performed with at least two biological replicates and several technical repeats.

### HBsAg measurements

For intracellular HBsAg analysis by FACS, cells were harvested by trypsinization and 1×10^5^ - 3×10^5^ cells/well were fixed with BD Cytofix/Cytoperm fixation/permeabilization solution (BD Biosciences) for 15 min at 4°C. Cells were washed twice with 1X BD Perm/Wash buffer (BD Biosciences) and stained with primary human HBsAg antibody (CR8097; US9512201 at 1.5 μg/ml) for 1h at 4°C. After two washes with BD Perm/Wash buffer, cells were stained with APC-conjugated mouse anti-human IgG antibody (Southern Biotech; 1:500) for ~45 min at 4°C. Following incubation, samples were washed with BD Perm/Wash buffer, resuspended in PBS and analyzed by BD FACSCanto II flow cytometer (BD Biosciences). Data were analyzed using FlowJo software (Treestar, Inc.).

HBsAg in culture medium was detected with an AlphaLISA developed in house. Five days post drug or siRNA treatment, supernatant from the culture plate was transferred to 384-well tissue-culture treated white opaque plate (Greiner Bio-one) and mixed with 50mM Tris assay buffer containing AlphaLisa acceptor and donor custom conjugated beads with in house generated human anti-HBsAg antibodies. Four hours after incubation at room temperature, the release of singlet oxygen was excited at 680nM and the emission of light signal was detected at 615nM using the Envision Plate Reader (Perkin Elmer).

For HBsAg analysis by immunofluorescence, cells were fixed with 4% (wt/vol) paraformaldehyde in PBS at room temperature for 15min. After washing with PBS, cells were incubated with blocking buffer [3% (wt/vol) BSA, 0.5% Triton X-100, 10% (vol/vol) FBS in 1× PBS] for 30-60 min followed by an hour incubation at room temperature with the HBsAg antibody (CR8097) diluted to 1.5 μg/mL in incubation buffer [3% (wt/vol) BSA, 0.5% Triton X-100 in 1× PBS]. The cells were washed twice with PBS and stained in the dark at room temperature for 30 min with NucBlue Fixed Cell ReadyProbes reagent - DAPI (ThermoFisher) and the secondary APC-conjugated mouse anti-human IgG antibody (Southern Biotech) diluted 1:1,000 in incubation buffer. After two washes in PBS, images were acquired with ImageXpress® and analyzed with MetaXpress®. For each condition, infections were done in triplicate with nine images acquired per replicate.

### Total intracellular HBV DNA analysis by qPCR

For total intracellular HBV DNA quantification, normalized total DNA extracted with DNeasy Blood & Tissue Kit (Qiagen) following manufacturer’s instructions was quantified by qPCR with primers described in (Zhu et al., 2011) using QuantiFast Probe PCR Master Mix (Qiagen). qPCR was run on ABI7900HT (Applied Biosystems) with PCR cycling conditions as follows: 95°C for 3min; and 95°C for 3s, 60°C for 30s for 40 cycles.

### RNA extraction and cDNA synthesis

RNA was extracted with TRIZOL (ThermoFisher) following manufacturer’s instructions. For cDNA synthesis, 500ng of total RNA was reverse transcribed using SuperScript VILO Master Mix at 42°C for 1 h with reverse transcriptase heat inactivated at 85°C for 5 min. The resulting cDNA was diluted 1:10 in water, and 1μL was used for qPCR analysis using FAST SYBR green master mix (Applied Biosystems) with 1X PrimeTime IDT primers. Each primer pair reaction was run in triplicate following on ABI7900HT (Applied Biosystems) with PCR cycling conditions as follows: 95°C for 20s; and 95°C for 3s, 60°C for 30s for 40 cycles.

### Northern blotting

To detect HBV transcripts, Northern blotting was executed as described in (Tropberger et al., 2015). Briefly, one microgram of RNA samples and 2μL of Millennium RNA ladder (ThermoFisher) were run on a denaturing 1% (wt/vol) agarose gel electrophoresis with reagents from the Northern Max Kit (Ambion) and transferred to Nytran membrane (TurboBlot kit) by using the TurboBlotter system (GE Healthcare). Ribosomal RNAs was visualized with methylene blue solution (Sigma-Aldrich). A DIG-labeled (−) strand HBV RNA probe was transcribed from ScaI-linearized pGEM3Z-HBV plasmid with DIG Northern Starter Kit (Roche) according to manufacturer’s instructions. Hybridization at 68°C, washes, and detection with CDP-star (Roche) were carried out according to manufacturer’s instructions. Images were acquired with the Azure c600 Imaging System (Azure Biosystems).

### RNA tail-length assay

RNA tail length was analyzed using the Poly(A) Tail-Length Assay Kit (ThermoFisher) following manufacturer’s instructions. Briefly, 1μg of RNA was incubated with Tail Buffer Mix and Tail Enzyme Mix at 37°C for 60min, and the reaction with stopped with 2μL of 10X Tail stop mix. 5μL of the resulting G/I tailed RNA was reverse transcribed and PCR amplified with HBV-specific Forward (ACCGACCTTGAGGCATACTT) and Reverse (ATAAGGGTCGATGTCCATGC) primers as a control, HBV-specific Forward and universal reverse primers for detecting the HBV specific RNA tail, or with the actin-specific forward and universal reverse primer supplied with the kit. PCR amplification was performed using following conditions: 94°C for 2min; 94°C for 10s and 60°C for 60s for 35 cycles; 72°C for 5min. A portion (12.5μL) of each PCR reaction was run on a 2% E-Gel Agarose Gel with ethidium bromide and images were acquired with the Azure c600 Imaging System (Azure Biosystems).

### TENT4B cloning

Human TENT4B was cloned as follows. Protein sequence of TENT4B was obtained from UniProtKB (accession Q8NDF8, Isoform 1, 572 amino acids). It was back translated into DNA sequence optimized for expression in *Spodoptera frugiperda* and synthesized by GeneArt Strings (Thermo Scientific) in a few fragments. The DNA fragments were cloned into pET30b vector by homologous recombination, so that the gene expression product is a fusion protein with an N-terminal hexahistidine tag followed by ZZ protein and 3C protease recognition site.

### TENT4B expression and purification

*E. coli* BL21(DE3) cells transformed by the plasmid was grown in 0.5 L auto-induction medium (Terrific Broth supplemented with 100mM sodium phosphate pH 7.0 buffer, 2mM MgSO_4_, trace metals, 0.05% glucose, 0.05% lactose and 100 μg/mL Kanamycin) at 37°C for 2 hours, then the temperature was reduced to 18°C and the cells were grown overnight. The harvested cells were resuspended in buffer A (25 mM HEPES pH 7.5, 300mM KCl, 5% glycerol, 20mM Imidazole, and 1mM TCEP), lysed by sonication, centrifuged at 50,000xG for 30 min. Supernatant was taken and incubated with Nickel Sepharose 6 Fast Flow (GE Healthcare) equilibrated with buffer A at 4°C for 2 hours. The slurry was loaded in a gravity column, washed with buffer A, then the bound protein was eluted with buffer B (same as buffer A except that Imidazole concentration was 500mM). Fractions containing TENT4B protein was collected and treated with 3C protease overnight, while dialyzed against buffer A without Imidazole. The protein solution was diluted with buffer C (25mM HEPES pH 7.5, 5% glycerol, and 1mM TCEP) to reduce the KCl concentration to less than 200mM, then loaded onto HiTrap Heparin column (GE Healthcare) equilibrated with buffer D (25mM HEPES pH 7.5, 200mM KCl, 5% glycerol, and 1mM TCEP). After column wash with buffer D, the protein was eluted with 0 to 100% gradient of buffer E (25mM HEPES pH 7.5, 1M KCl, 5% glycerol, and 1mM TCEP) over 20 column volumes. The fractions containing TENT4B was pooled and concentrated, then loaded onto Superdex 75 26/60 column (GE Healthcare) at a flow rate of 2.5 ml/min. The fractions containing TENT4B was pooled and concentrated, however due to low solubility of the protein, the final concentration was 0.19 mg/ml. The protein was aliquoted, flash frozen, and stored at −80°C.

### TENT4B ATP depletion assay

The TENT4B ATP depletion activity was performed in assay buffer (25mM Tris-HCL pH 7.5, 50mM KCl, 5mM MgCl2, 325mM EDTA, 0.02% Tween-20, 5% glycerol, 1mM DTT, 0.01% BSA, 2% DMSO) containing 15nM purified TENT4B, 125nM in poly(A_15_) and 1μM ATP. ATP depletion was readout at multiple time-points using Kinase Glo kit following manufacturer’s instructions (Promega).

### Co-immunoprecipitation

HepG2-HBV cells were washed with PBS and lysed in Pierce™ IP lysis buffer (ThermoFisher) with complete protease inhibitor (Roche) + phosphatase inhibitor (Roche). To immunoprecipitate protein complexes, Dynabeads™ Protein G (Invitrogen) were used according to manufacturer’s instructions. Briefly, cleared cell lysates were incubated with rotation overnight at 4°C with Dynabeads™ Protein G pre-bound with 1-5μg of anti-PAPD5 (Sigma-Aldrich; HPA042968), anti-ZCCHC14 (Novus; NBP1-71852) or 5μg IgG control (Millipore; 12-370) antibodies. Following immunoprecipitation, the Dynabeads®-Ab-Ag complex was washed 3 times using 200μL IP Lysis buffer. To avoid co-elution of proteins bound to the tube wall, the Dynabeads®-Ab-Ag complex was transferred to a new tube after the final wash. To elute target antigen, the Dynabeads®-Ab-Ag was resuspended in 1X NuPage LDS sample buffer and NuPage Sample Reducing Agent and heated for 10 min at 70°C. Immunoprecipitated samples and the 2% input samples were loaded onto a 4-12% Bis-Tris gel and analyzed by western blotting.

### Cellular thermal shift assay

HepG2-HBV cells were treated with 10μM RG7834 or DMSO for 1h at 37°C in PBS containing protease inhibitor before being heated for 3 min at various temperatures. Cells were snap frozen in liquid nitrogen and lysed by repeated freeze thaw cycle. Cell lysates were cleared by centrifugation to remove precipitated and aggregated proteins. Protein expression levels were determined by western blotting and quantified by densitometry.

### Western blotting

Cultured cells were washed with PBS and lysed in RIPA lysis buffer (Boston Bioproducts) with complete protease inhibitor (Roche) + phosphatase inhibitor (Roche). Clarified cell extracts were boiled in1X NuPage LDS sample buffer (Invitrogen; NP0007) and 1X NuPage Sample Reducing Agent (Invitrogen; NP0009). Equal protein extracts were loaded onto a 4-12% Bis-Tris gel and transferred to nitrocellulose membranes using standard wet electroblotting system (Bio-Rad). Membranes were blocked in Odyssey blocking buffer (LI-COR Biosciences, Lincoln, NE, USA), and proteins of interest were analyzed using primary antibodies as follows: anti-PAPD5 (1:500; Sigma-Aldrich; HPA042968), anti-ZCCHC14 (1:2000; Novus; NBP1-71852) and anti-tubulin (1:3000; Invitrogen; MA5-16308). Secondary antibodies were used IRDye 680-conjugated or 800-conjugated donkey anti-mouse or goat anti-rabbit antibodies (1:10,000; LI-COR Biosciences). Signal intensities were quantified using Image Studio™ Lite quantification software.

## SUPPLEMENTAL INFORMATION TITLES

**Figure S1.**
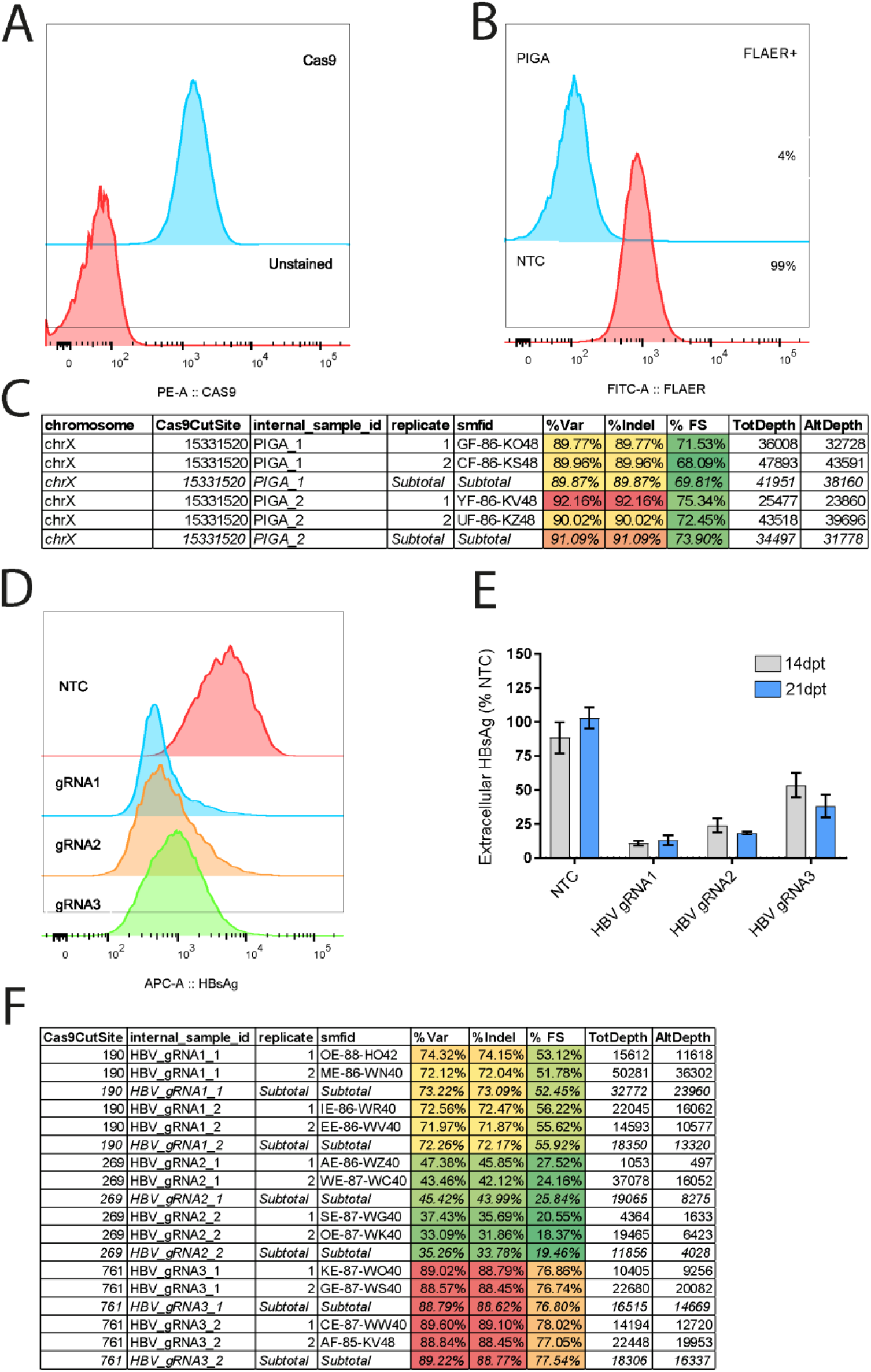
Generation and validation of the Cas9 HepG2-HBV cell line for genome-wide CRISPR screen. **(A)** FACS analysis of Cas9 expression in Cas9 HepG2-HBV cell line. **(B)** FACS-based FLAER analysis of Cas9 editing using sgRNA targeting PIG-A enzyme at 21 days post-transduction (dpt). **(C)** On-target NGS analysis of Cas9 editing across the PIG-A gene locus at 14 dpt. **(D-E)** FACS analysis of intracellular HBsAg levels at 14 dpt **(D)** and AlphaLISA analysis of extracellular HBsAg levels **(E)** in Cas9 HepG2-HBV cells transduced with lentiviruses expressing a NTC sgRNA and three sgRNAs targeting the HBV genome. **(F)** On-target NGS analysis of Cas9 editing across the HBV genome at 14 dpt.

**Figure S2.**
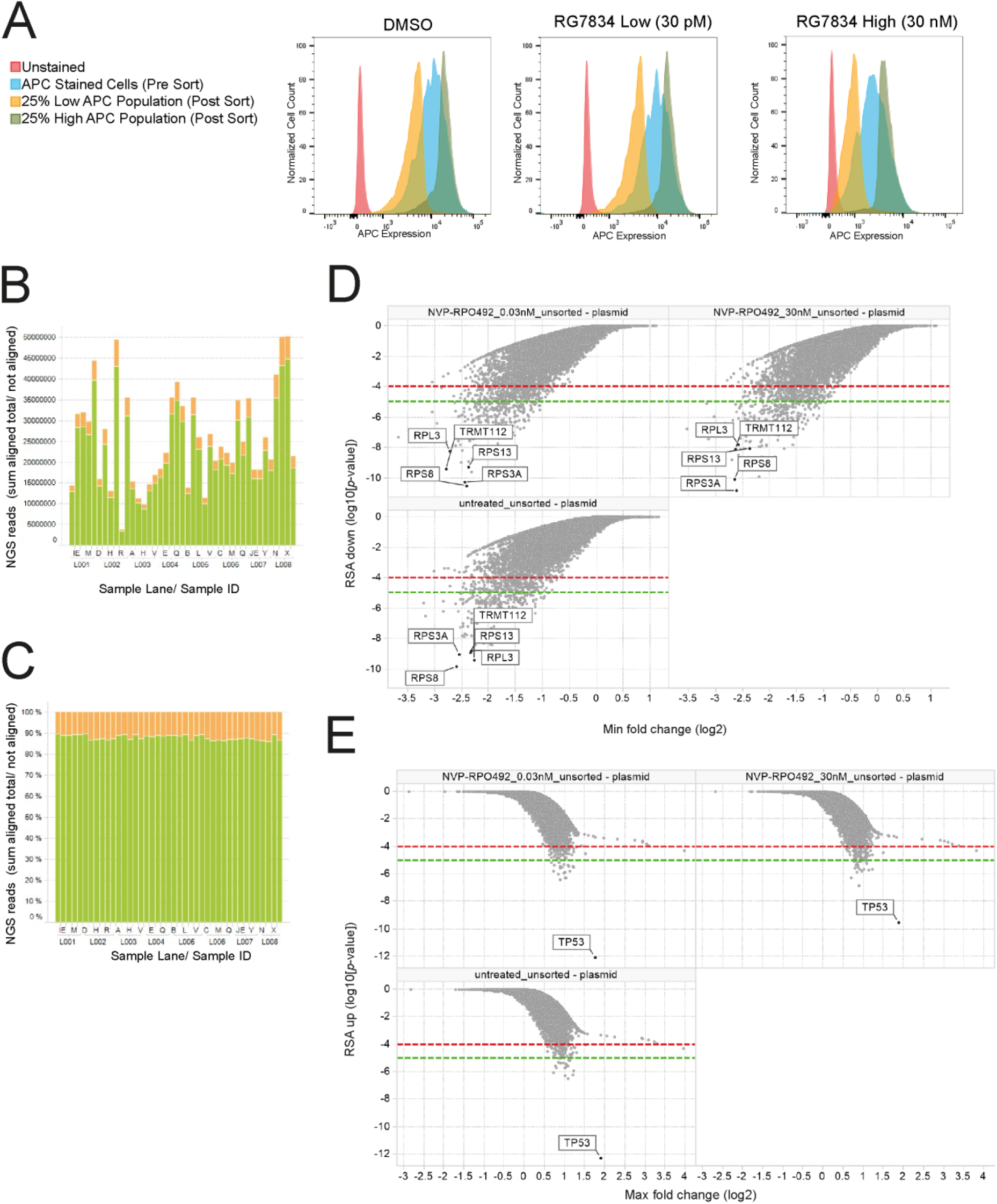
Genome-wide CRISPR screen analysis. **(A)** FACS analysis of Cas9 HepG2-HBV cells selected following pooled lentiviral sgRNA library transduction and treatment with DMSO, RG7834 low dose (30pM) or high dose (30nM). Cells were stained using an antibody for HBsAg and sorted by FACS into HBsAg-high and HBsAg-low populations. **(B)** NGS analysis of the total reads per sample. **(C)** Analysis of the alignment of the NGS reads to expected barcodes from the pooled libraries (green: sum of aligned reads to expected barcodes; orange: sum of not aligned reads). **(D-E)** Genome-wide CRISPR screen analysis on the sgRNA counts in unsorted samples compared to the sgRNA counts in the input CRISPR library. **(D)** Waterfall plot of sgRNAs found in lower than expected abundance in the unsorted populations. **(E)** Waterfall plot of sgRNAs found in higher than expected abundance in the unsorted populations.

**Figure S3.**
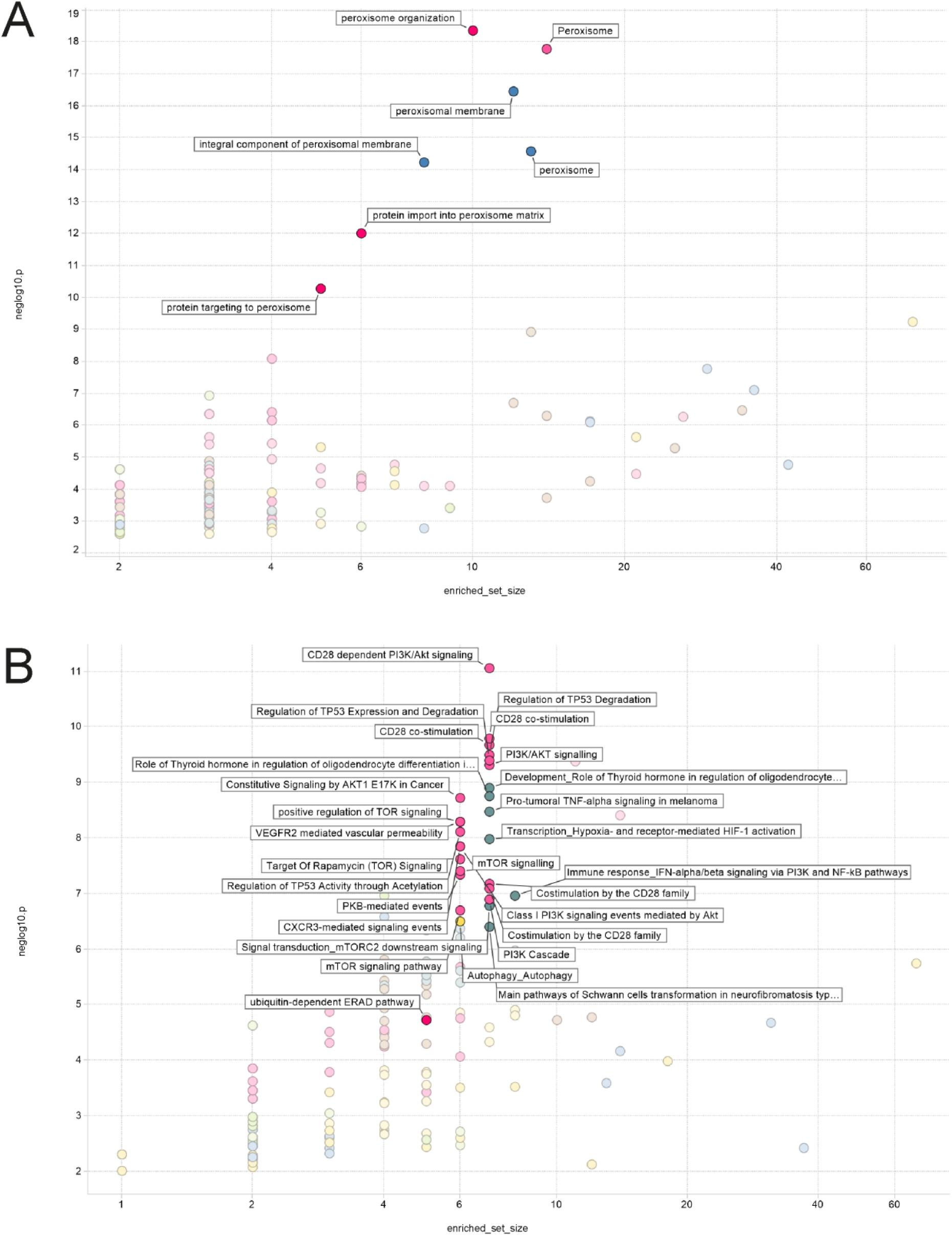
Enrichment pathway analysis. Enrichment pathway analysis of **(A)** pro-HBsAg genes and **(B)** anti-HBsAg genes.

**Figure S4.**
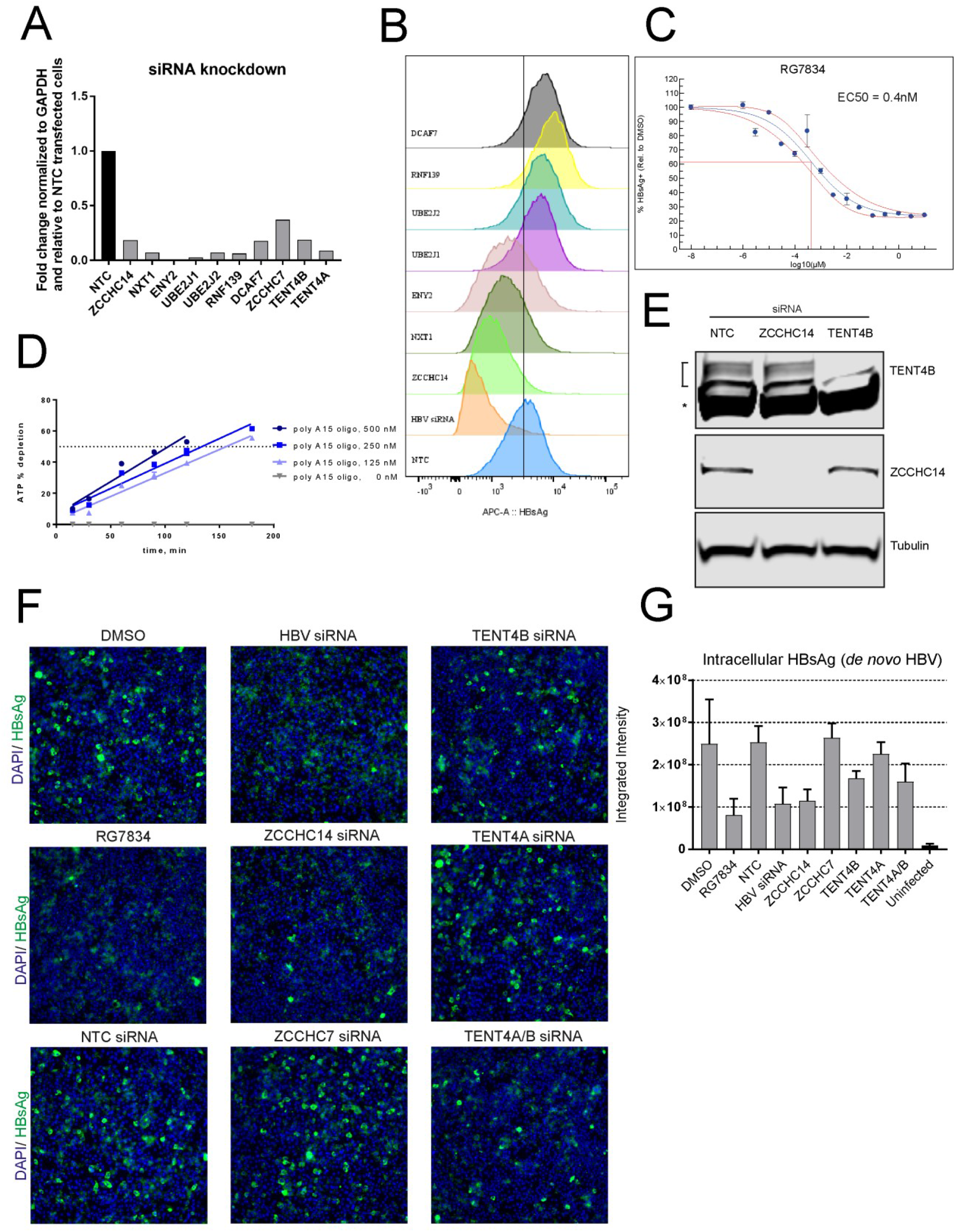
Validation of host factors required for HBsAg production. **(A)** siRNA-mediated gene knockdown confirmed by qPCR. **(B)** FACS analysis of intracellular HBsAg levels following siRNA-mediated knockdown of the putative pro-HBsAg and anti-HBsAg genes in HepG2-HBV for 5 days. The graph shows representative FACS plots of two biological replicates. **(C)** EC50 curve analysis of RG7834 inhibition of intracellular HBsAg expression measured by FACS following 10 days treatment. **(D)** Kinase-Glo (Promega) analysis of RNA-dependent ATP depletion by purified TENT4B. **(E)** Western blot analysis of TENT4B and ZCCHC14 protein levels following siRNA-mediated gene knockdown for 6 days in HepG2-HBV cells. A bracket marks several TENT4B isoforms. *, in TENT4B blots indicates a non-specific band. **(F)** Immunofluorescence analysis of intracellular HBsAg at 7 days post de novo infection with HBV in HepG2-NTCP1 cells. Representative images of two experiments are shown. **(G)** Immunofluorescence analysis of integrated intensity of intracellular HBsAg signal using MetaXpress. Results of at least two independent replicates are shown. Error bars represent mean ± SD.

**Table S1.** Peroxisome-associated genes involved in HBsAg regulation identified by genome-wide CRISPR screen.

| Gene ID | Symbol | Description | GO process |
| --- | --- | --- | --- |
| 215 | ABCD1 | ATP binding cassette subfamily D member 1 | alpha-linolenic acid metabolic process;fatty acid beta-oxidation using acyl-CoA oxidase;peroxisomal long-chain fatty acid import;peroxisomal membrane transport;peroxisome organization |
| 51 | ACOX1 | acyl-CoA oxidase 1 | alpha-linolenic acid metabolic process;fatty acid beta-oxidation using acyl-CoA dehydrogenase;fatty acid beta-oxidation using acyl-CoA oxidase; generation of precursor metabolites and energy;lipid homeostasis;lipid metabolic process;prostaglandin metabolic process;protein targeting to peroxisome |
| 3295 | HSD17B4 | hydroxysteroid 17-beta dehydrogenase 4 | alpha-linolenic acid metabolic process;androgen metabolic process;bile acid biosynthetic process;estrogen metabolic process;fatty acid beta-oxidation using acyl-CoA oxidase;medium-chain fatty-acyl-CoA metabolic process;protein targeting to peroxisome |
| 5189 | PEX1 | peroxisomal biogenesis factor 1 | microtubule-based peroxisome localization;peroxisome organization;protein import into peroxisome matrix;protein targeting to peroxisome |
| 5192 | PEX10 | peroxisomal biogenesis factor 10 | peroxisome organization;protein import into peroxisome matrix;protein targeting to peroxisome;protein ubiquitination |
| 5193 | PEX12 | peroxisomal biogenesis factor 12 | peroxisome organization;protein import into peroxisome matrix;protein monoubiquitination;protein targeting to peroxisome;protein ubiquitination |
| 5194 | PEX13 | peroxisomal biogenesis factor 13 | cerebral cortex cell migration;fatty acid alpha-oxidation;locomotory behavior;microtubule-based peroxisome localization;neuron migration;protein import into peroxisome matrix, docking; protein ubiquitination |
| 9409 | PEX16 | peroxisomal biogenesis factor 16 | ER-dependent peroxisome localization;ER-dependent peroxisome organization;peroxisome membrane biogenesis;peroxisome organization;protein import into peroxisome matrix;protein import into peroxisome membrane; protein to membrane docking |
| 5824 | PEX19 | peroxisomal biogenesis factor 19 | chaperone-mediated protein folding and transport;establishment of protein localization to peroxisome;negative regulation of lipid binding;peroxisome fission;peroxisome organization;protein import into peroxisome membrane;protein stabilization; transmembrane transport |
| 5828 | PEX2 | peroxisomal biogenesis factor 2 | fatty acid beta-oxidation; negative regulation of transcription by RNA polymerase II;peroxisome organization;protein destabilization;protein import into peroxisome matrix;protein targeting to peroxisome;protein ubiquitination;very long-chain fatty acid metabolic process |
| 55670 | PEX26 | peroxisomal biogenesis factor 26 | protein import into peroxisome matrix;protein import into peroxisome membrane;protein targeting to peroxisome |
| 8504 | PEX3 | peroxisomal biogenesis factor 3 | peroxisome membrane biogenesis;peroxisome organization;protein import into peroxisome membrane;transmembrane transport |
| 5190 | PEX6 | peroxisomal biogenesis factor 6 | peroxisome organization;protein import into peroxisome matrix;protein stabilization;protein targeting to peroxisome |
| 6342 | SCP2 | sterol carrier protein 2 | alpha-linolenic acid metabolic process;bile acid biosynthetic process;fatty acid beta-oxidation using acyl-CoA oxidase;lipid hydroperoxide transport;phospholipid transport;positive regulation of intracellular cholesterol transport;protein targeting to peroxisome;steroid biosynthetic process;sterol transport |

